# Sensitisation of colonic nociceptors by TNFα is dependent on TNFR1 expression and p38 MAPK activity

**DOI:** 10.1101/2022.02.06.478183

**Authors:** Katie H. Barker, James P. Higham, Luke A. Pattison, Toni S. Taylor, Iain P. Chessell, Fraser Welsh, Ewan St. J. Smith, David C. Bulmer

## Abstract

Visceral pain is a leading cause of morbidity in gastrointestinal diseases, which is exacerbated by the gut related side-effects of many analgesics. New treatments are needed and further understanding of the mediators and mechanisms underpinning visceral nociception in disease states is required to facilitate this. The pro-inflammatory cytokine TNFα is linked to pain in both patients with inflammatory bowel disease and irritable bowel syndrome, and has been shown to sensitise colonic sensory neurons. Somatic, TNFα- triggered thermal and mechanical hypersensitivity is mediated by TRPV1 signalling and p38 MAPK activity respectively, downstream of TNFR1 receptor activation. We therefore hypothesised that TNFR1-evoked p38 MAPK activity may also be responsible for TNFα sensitisation of colonic afferent responses to the TRPV1 agonist capsaicin, and noxious distension of the bowel. Using Ca^2+^ imaging of dorsal root ganglion sensory neurons, we observed TNFα-mediated increases in intracellular [Ca^2+^] and sensitisation of capsaicin responses. The sensitising effects of TNFα were dependent on TNFR1 expression and attenuated by p38 MAPK inhibition. Consistent with these findings, *ex vivo* colonic afferent fibre recordings demonstrated enhanced response to noxious ramp distention of the bowel and bath application of capsaicin following TNFα pre-treatment. Responses were reversed by p38 MAPK inhibition and absent in tissue from TNFR1 knockout mice. Our findings demonstrate a contribution of TNFR1, p38 MAPK and TRPV1 to TNFα-induced sensitisation of colonic afferents, highlighting the potential utility of these drug targets for the treatment of visceral pain in GI disease.

**Abstract figure legend:** TNFα sensitised Ca^2+^ responses to the TRPV1 agonist capsaicin in dorsal root ganglion sensory neurons. Sensitisation was TNFR1-dependent and attenuated by inhibition of p38 MAPK. Direct Ca^2+^ responses to TNFα were TRPV1-and TRPA1-dependent. In *ex vivo* colonic afferent recordings, TNFα increased sensitivity to noxious ramp distension and capsaicin, both of which were absent in TNFR1^-/-^ tissue or blocked by inhibition of p38 MAPK. These findings establish a role for TNFR1, p38 MAPK and TRPV1 in TNFα-mediated sensitisation of colonic afferents.

**Key Points Summary:**

- TNFα sensitises sensory neurons and colonic afferents to the TRPV1 agonist capsaicin.
- TNFα-mediated sensitisation of sensory neurons and colonic nociceptors is dependent on TNFR1 expression.
- TNFα sensitisation of sensory neurons and colonic afferents to capsaicin and noxious ramp distension is abolished by inhibition of p38 MAPK.

## Introduction

Abdominal pain is a leading cause of morbidity in gastrointestinal (GI) diseases, such as inflammatory bowel disease (IBD), and irritable bowel syndrome (IBS).(Spiller & Major, 2016; Grundy *et al*., 2019). Mechanistically, the sensitisation of visceral nociceptors by inflammation, infection, allergy, and other pathological processes, is a principal cause of pain in disease states, although the mediators and mechanisms underpinning this are not yet fully understood (McMahon *et al*., 2015; Grundy *et al*., 2019). The proinflammatory cytokine tumour necrosis factor α (TNFα) has been implicated in the development of peripheral sensitisation and visceral pain in IBD and IBS patients, based on its causative role in inflammatory disease pathology, localised release from mast cells, and the significant correlation between pain scores and peripheral blood mononuclear cell (PBMC)-evoked TNFα release in IBS patients (Rijnierse *et al*., 2006; Hughes *et al*., 2013). These observations are supported by expression of the TNFR1 receptor in mouse colonic sensory neurons (Hockley *et al*., 2019) and human dorsal root ganglion (DRG) neurons (Wangzhou *et al*., 2020), as well as functional data demonstrating the sensitisation of colonic neurons and colonic afferents by TNFα (Hughes *et al*., 2013). These effects have been linked to enhanced tetrodotoxin (TTX)-resistant voltage gated sodium channel currents (Na_V_1.8 subtype) (Jin & Gereau IV, 2006) and suppressed voltage gated potassium channel currents (delayed rectifier subtypes), downstream of TNFR1 receptor activation in DRG neurons (Ibeakanma & Vanner, 2010), as well as there being a role for TRPA1 channel activity in colonic afferents ((Hughes *et al*., 2013). The translational importance of these findings is supported by studies utilising IBD patient colonic biopsy supernatants or IBS patient PBMCs, which have confirmed the essential contribution of TNFα to the respective sensitisation of DRG neurons or colonic afferents using tissue from TNFR1^-/-^ mice or pre-treatment with the anti-TNFα monoclonal antibody infliximab (Ibeakanma & Vanner, 2010; Hughes *et al*., 2013).

In addition to visceral pain, TNFα also evokes somatic, thermal, and mechanical hypersensitivity by increasing TRPV1 activity (Khan *et al*., 2008) and p38 mitogen-activated protein kinase (MAPK)-mediated Na_V_1.8 activity, downstream of TNFR1 receptor activation (Jin & Gereau IV, 2006). Given TRPV1 and Na_V_1.8 channels are co-expressed with TNFR1 in colonic nociceptors (Hockley *et al*., 2019), we reasoned that TNFα may sensitise responses to the TRPV1 agonist capsaicin, and that p38 MAPK signalling may be responsible for TNFα-mediated sensitisation of colonic nociceptors. Consequently, the aims of this study were to investigate the contribution of p38 MAPK and TNFR1 to TNFα-mediated sensitisation of colonic afferents to capsaicin and noxious distension of the bowel using a combination of Ca^2+^ imaging of DRG neurons and ex vivo electrophysiological recordings of colonic afferent activity.

## Materials and Methods

### Ethical Approval

All animal experiments were conducted in compliance with the Animals (Scientific Procedures) Act 1986 Amendment Regulations 2012 under Project Licence P7EBFC1B1 granted to E. St. J. Smith by the Home Office and approval by the University of Cambridge Animal Welfare Ethical Review Body.

### Reagents

Stock concentrations of TNFα (0.1 mg/ml; H_2_O with 0.2% (w/v) bovine serum albumin), capsaicin (1 mM; 100% ethanol) and staurosporine (10 mM; DMSO) were dissolved as described, all purchased from Sigma-Aldrich. R7050 (10 mM; DMSO), thapsigargin (1 mM; DMSO), A425619 (1 mM; DMSO) and SB203580 (10 mM; DMSO) were obtained from Tocris and stock concentrations made up as described. Nifedipine (100 mM; DMSO) and atropine (100 mM; 100% ethanol) were purchased from Sigma-Aldrich and dissolved as described. All drugs were diluted to working concentrations in extracellular solution (ECS) or Krebs buffer on the day of the experiment.

### Animals

Adult male C57BL/6J mice (8-16 weeks) were obtained from Charles River (Cambs, UK; RRID:IMSR_JAX:000664). Mice were conventionally housed in temperature-controlled rooms (21°C) with a 12-h light/dark cycle and provided with nesting material, a red plastic shelter and access to food and water ad libitum. TNFR1^-/-^ mice (9-14 weeks; 8F, 7M; Jackson Laboratory, ME, USA; RRID:IMSR_JAX:003242) were housed in individually ventilated plastic cages under the same conditions.

### Primary culture of mouse dorsal root ganglion neurons

DRG neurons were cultured as previously described (Hockley *et al*., 2019). Briefly, mice were euthanised by exposure to a rising concentration of CO_2_, followed by cervical dislocation. Isolated DRG (T12-L5, spinal segments innervating the distal colon) were incubated in 1 mg/ml collagenase (15 min) followed by trypsin (1 mg/ml) both with 6 mg/ml BSA in Leibovitz’s L-15 Medium, GlutaMAX™ Supplement (supplemented with 2.6% (v/v) NaHCO_3_). DRG were resuspended in 2 ml Leibovitz’s L-15 Medium, GlutaMAX™ Supplement containing 10% (v/v) foetal bovine serum (FBS), 2.6% (v/v) NaHCO_3_, 1.5% (v/v) glucose and 300 units/ml penicillin and 0.3 mg/ml streptomycin (P/S). DRG were mechanically dissociated, centrifuged (1000 rpm) and the supernatant collected for five triturations. Following centrifugation and resuspension, the supernatant (50 μL) was plated onto 35 mm poly-D-lysine coated glass bottom culture dishes (MatTek, MA, USA), and further coated with laminin (Thermo Fisher: 23017015). Dishes were incubated for 3 hours to allow cell adhesion, before flooding with 2 ml Leibovitz’s L-15 Medium, GlutaMAX™ Supplement containing 10% (v/v) FBS, 2.6% (v/v) NaHCO_3_, 1.5% (v/v) glucose and P/S and cultured for 24 hours. All incubations were carried out at 37 °C with 5% CO_2_.

### Ca^2+^ imaging

Extracellular solution (in mM: 140 NaCl, 4 KCl, 1 MgCl_2_, 2 CaCl_2_, 4 glucose, 10 HEPES) was prepared and adjusted to pH 7.4 using NaOH and an osmolality of 290-310 mOsm using sucrose. Cells were incubated for 30 min with 100 μl of 10 μM Ca^2+^ indicator Fluo-4-AM (room temperature; shielded from light). For inhibitor studies requiring pre-incubation, 200 μl of drug was added for 10 min prior to imaging.

Dishes were mounted on the stage of an inverted microscope (Nikon Eclipse TE-2000S) and cells were visualised at 10 x magnification with brightfield illumination. Cells were initially superfused with ECS, or drug in inhibitor studies to establish baseline. For studies in which Ca^2+^ was absent from ECS, bath solution was made up as follows (in mM): 140 NaCl, 4 KCl, 2 MgCl_2_, 4 glucose, 10 HEPES, 1 EGTA (pH 7.35-7.45 with NaOH; 290-310 mOsm with sucrose). To compensate for the loss of extracellular divalent cations, the MgCl_2_ concentration was increased and EGTA was used to chelate any remaining Ca^2+^.

Fluorescent images were captured with a CCD camera (Rolera Thunder, Qimaging, MC, Canada or Retiga Electro, Photometrics, AZ, USA) at 2.5 fps with 100 ms exposure and a 470 nm light source for excitation of Fluo-4-AM (Cairn Research, Faversham, UK). Emission at 520 nm was recorded with μManager (Edelstein *et al*., 2014). All protocols began with a 10 s baseline of ECS before drug superfusion. With multiple drug additions to the same dish, cells were allowed 4 min recovery between applications. Finally, cells were stimulated with 50 mM KCl for 10 s to determine cell viability, identify neuronal cells and allow normalisation of fluorescence. A fresh dish was used for each protocol and all solutions were diluted in ECS.

### Ca^2+^ imaging data analysis

Individual cells were circled on a brightfield image and outlines overlaid onto fluorescent images using ImageJ (NIH, MA, USA). Pixel intensity was measured and analysed with custom-written scripts in RStudio (RStudio, MA, USA). Background fluorescence was subtracted from each cell, and fluorescence intensity (F) baseline corrected and normalised to the maximum fluorescence elicited during 50 mM KCl stimulation (F_pos_). Maximum KCl fluorescence was denoted as 1 F/F_pos_. Further analysis was confined to cells with a fluorescence increase ≥ 5 standard deviations above the mean baseline before 50 mM KCl application. Neurons were deemed responsive to a drug challenge if a fluorescence increase of 0.1 F/F_pos_ was seen in response to drug perfusion. The proportion of responsive neurons and magnitude of the fluorescence response was measured for each experiment, with peak responses calculated from averaging fluorescence values of individual neurons at each time point.

### Magnetic-activated cell sorting

To determine the role of satellite cells, such as glia, in the neuronal responses to TNFα, satellite cells were removed from DRG cultures using magnetic-activated cell sorting (MACS), with equipment purchased from Miltenyi Biotec and using protocols previously described (Thakur *et al*., 2014).

DRG from 2-3 mice were isolated and cultured as above, but trypsin incubation was omitted and DRG were incubated with collagenase (1 mg/ml with 6 mg/ml BSA) for 45 min. Pelleted neurons were washed in 2 ml Dulbecco’s phosphate-buffered saline (DPBS, containing 0.9 mM CaCl_2_ and 0.5 mM MgCl_2_) and centrifuged for 7 min (1000 rpm). The pellet was resuspended in MACS rinsing solution (120 μl), supplemented with 0.5% w/v BSA (sterile filtered at 0.2 μM), and incubated (5 min at 4°C) with a biotin-conjugated non-neuronal antibody cocktail (30 μl). DPBS was added to a volume of 2 ml and the suspension centrifuged for 7 min at 1000 rpm. The pellet was resuspended in 120 μl MACS rinsing solution with 30 μL biotin-binding magnetic beads and incubated for a further 10 min at 4 °C, before being topped-up to 500 μl with MACS buffer.

The cell suspension was filtered by gravity through a magnetic column (LD column), primed with 2.5 ml MACS rinsing solution. Following the addition of the cell suspension, 1 ml MACS rinsing solution was used to collect the remnants of the cell suspension and passed through the column prior to a final wash. The 5 ml elute was centrifuged for 7 min at 1000 rpm and the final pellet resuspended in supplemented L-15 medium, before plating on 35 mm poly-D-lysine coated glass bottom culture dishes further coated with Matrigel (diluted 1:10 in L-15 medium). Dishes were incubated for 3 hours to allow cell adhesion, after which 2 ml Leibovitz’s L-15 Medium, GlutaMAX™ Supplement containing 10% (v/v) FBS, 2.6% (v/v) NaHCO_3_, 1.5% (v/v) glucose and P/S was added, and dishes were cultured for 48 hours (37°C, 5% CO_2_). Medium was changed after 24 hours.

### Immunocytochemistry of cultured DRG

DRG neurons were cultured as above and seeded onto 12 mm coverslips coated in poly-L-lysine and laminin. After 24-48 hours in culture, cells were fixed at room temperature in 4%, pH 7.0 paraformaldehyde (10 min) and washed in PBS. Cells were permeabilized with 0.05% Triton-X100 for 5 min at room temperature. Cells were washed again in PBS and then blocking buffer (1% goat serum in 0.2% Triton-X100) was applied. After 2 hours, cells were incubated with a rabbit anti-βIII-tubulin primary antibody (1:1000, Abcam: ab18207; RRID:AB_444319) (Prado *et al*., 2021) for 3 hours at room temperature.

Following primary antibody incubation, cells were washed in PBS and incubated with an Alexa Fluor-568 goat anti-rabbit secondary antibody diluted in PBS (1:1000, Invitrogen: A11008; RRID:AB_143165) (Crerar *et al*., 2019) plus 4’-6-diamidino-2-phenylindole (DAPI; 1:1000, Abcam) for 1 hour at room temperature. After a final wash, coverslips were mounted, cell side down, on 25 × 75 × 1 mm glass slides using Mowoil 4-88 mounting medium (Sigma-Aldrich: 81381). Mounting medium was set at 4°C and slides were imaged within 1 hour.

Slides were imaged using an Olympus BX51 microscope. Fluorophores were excited with 568 nm (Alexa Fluor-568) or 350 nm (DAPI) light sources. Images were captured on a Qicam CCD camera (QImaging) with a 100 ms exposure and false coloured (βIII-tubulin, green; DAPI, blue). No βIII-tubulin staining was observed when the primary antibody was omitted (data not shown).

### Image Analysis

Images were analysed using ImageJ as previously described (Hartig, 2013). An automatic ‘minimum error’ threshold algorithm was applied to 8-bit images of βIII-tubulin or DAPI staining to distinguish background from objects. Binary and raw images were manually compared, and the threshold manually adjusted to ensure all regions of interest were captured. The threshold was placed at the first minimum after the major peak of the image histogram. Binary images then underwent watershed segmentation to separate distinct objects in proximity. Identified particles, positive for either βIII-tubulin or DAPI, were automatically counted using ImageJ and a ratio of βIII-tubulin-positive cells (neurons) to DAPI-positive cells (neurons and satellite cells) calculated.

### Ex vivo electrophysiology recordings of colonic afferent activity

Conducted as previously described (Hockley *et al*., 2020), the distal colorectum and associated lumbar splanchnic nerve (LSN; rostral to inferior mesenteric ganglia) were isolated from mice euthanised as described above and cannulated in a rectangular recording chamber with Sylgard base (Dow Corning, UK). Colons were luminally perfused (200 μl/min) and serosally superfused (7 ml/min; 32-34 °C) with carboxygenated Krebs buffer solution (in mM: 124 NaCl, 4.8 KCl, 1.3 NaH_2_PO_4_·H_2_O, 2.5 CaCl_2_·2H_2_O, 1.2 MgSO_4_·7H_2_O, 11.1 D-(+)-glucose, and 25 NaHCO_3_) supplemented with 10 μM atropine and 10 μM nifedipine to paralyse smooth muscle activity (Ness & Gebhart, 1988*a*).

Multi-unit activity from LSN bundles were recorded using borosilicate glass suction electrodes, and signals were amplified, band pass filtered (gain 5 KHz; 100–1300 Hz; Neurolog, Digitimer Ltd, UK), and filtered digitally for 50 Hz noise (Humbug, Quest Scientific, Canada). Analogue signals were digitized at 20 kHz (Micro1401; Cambridge Electronic Design, UK). All signals were visualised using Spike2 software.

### Electrophysiology Protocols

Following a minimum 30 min stabilisation period, repeated ramp distensions (0-80 mmHg) of the colorectum were performed by occluding luminal perfusion out-flow of the cannulated tissue (total distension time taking approximately 220 s). Ramp distensions at pressures > 30mmHg are noxious evoking pain behaviours in mice and humans (Ness & Gebhart, 1988*b*; Hughes *et al*., 2009).

In total, five ramp distensions, were performed separated by 15 min, after which 1 μM capsaicin (20 ml) was applied by bath superfusion (15 min after the last distension). TNFα (100 nM) or vehicle (buffer) was applied by luminal perfusion TNFα (15 min) was applied between the end of ramps 3 and 4. For experiments examining the effect of p38 MAPK inhibition, preparations were pre-treated by luminal perfusion with either SB203580 (10 μM) or vehicle (0.01% DMSO) started 15 min prior to and continued throughout TNFα luminal perfusion.

### Electrophysiological Data Analysis

In electrophysiological recordings, nerve discharge was determined by measuring the number of spikes passing a manually determined threshold twice the level of background noise (typically 60-80 μV) and binned to determine average firing frequency every 10 s. Changes in neuronal firing rates were calculated by subtracting baseline firing (averaged 3 min prior to distension or drug perfusion) from increases in nerve activity following ramp distension or capsaicin application. Peak firing to noxious mechanical distension and capsaicin application was determined respectively as the highest neuronal activity during ramp distension 5 and during the 10 min post capsaicin application. Changes to neuronal activity were recorded with each 5 mmHg increase in pressure and used to visualise ramp profiles. Capsaicin response profiles were plotted from binned data at 30 s increments. The area under the curve (AUC) was calculated for the duration of each ramp distension (0-80 mmHg) and for the 10 min following initial capsaicin application from response profiles using GraphPad Prism 9 software.

### Statistical analysis

All data sets were normality tested with a Shapiro-Wilk test and analysed using the appropriate statistical tests. The level of significance was set at p ≤ 0.05. All data are displayed as means ± standard deviation (SD). For Ca^2+^ imaging analysis, n represents the total number of dishes and N represents the total number of mice from which they were cultured. In MACS cultures, N represents the total number of independent pooled cultures.

## Results

### TNFα sensitises TRPV1 signalling in DRG neurons

In keeping with studies showing TNFα potentiation of TRPV1-mediated currents and Ca^2+^ flux in sensory neurons (Khan *et al*., 2008; Hsu *et al*., 2017), we examined the effect of overnight (24 hours) incubation, or acute application of TNFα (3nM), on capsaicin-evoked increases in intracellular Ca^2+^ concentration ([Ca^2+^]_i_) within DRG sensory neurons. Overnight incubation with TNFα elicited a significantly greater peak increase in [Ca^2+^]_i_ to 10 s 1 μM capsaicin (p = 0.028, two-way ANOVA with Holm-Šídák’s multiple comparisons; n = 5, N = 5; Figure 1A & B) in a similar proportion of neurons compared with vehicle incubation (p = 0.547, two-way ANOVA with Holm-Šídák’s multiple comparisons; n = 5, N = 5; Figure 1C). No sensitisation was observed at lower concentrations of capsaicin (0.01 μM: p = 0.404; 0.1 μM: p = 0.404, two-way ANOVA with Holm-Šídák’s multiple comparisons; n = 5, N = 5) and the proportion of capsaicin-sensitive neurons was unchanged following incubation with TNFα (0.01 μM: p = 0.901; 0.1 μM: p = 0.901, two-way ANOVA with Holm-Šídák’s multiple comparisons; n = 5, N = 5). Additionally, acute administration of TNFα (1 min), between repeat applications of capsaicin, prevented the marked desensitisation of response to the second capsaicin application (Figure 1D). This effect was only observed in a subset of capsaicin responsive neurons (29.93 ± 11.76%) that were co-sensitive to TNFα and represented 82.15 ± 20.91% of total capsaicin responders (e.g., cap_2_/cap_1_ p = 0.002, p = 0.004, one-way ANOVA with Holm-Šídák’s multiple comparisons test; n = 5-6, N = 5-6; Figure 1E).

**Figure 1.**
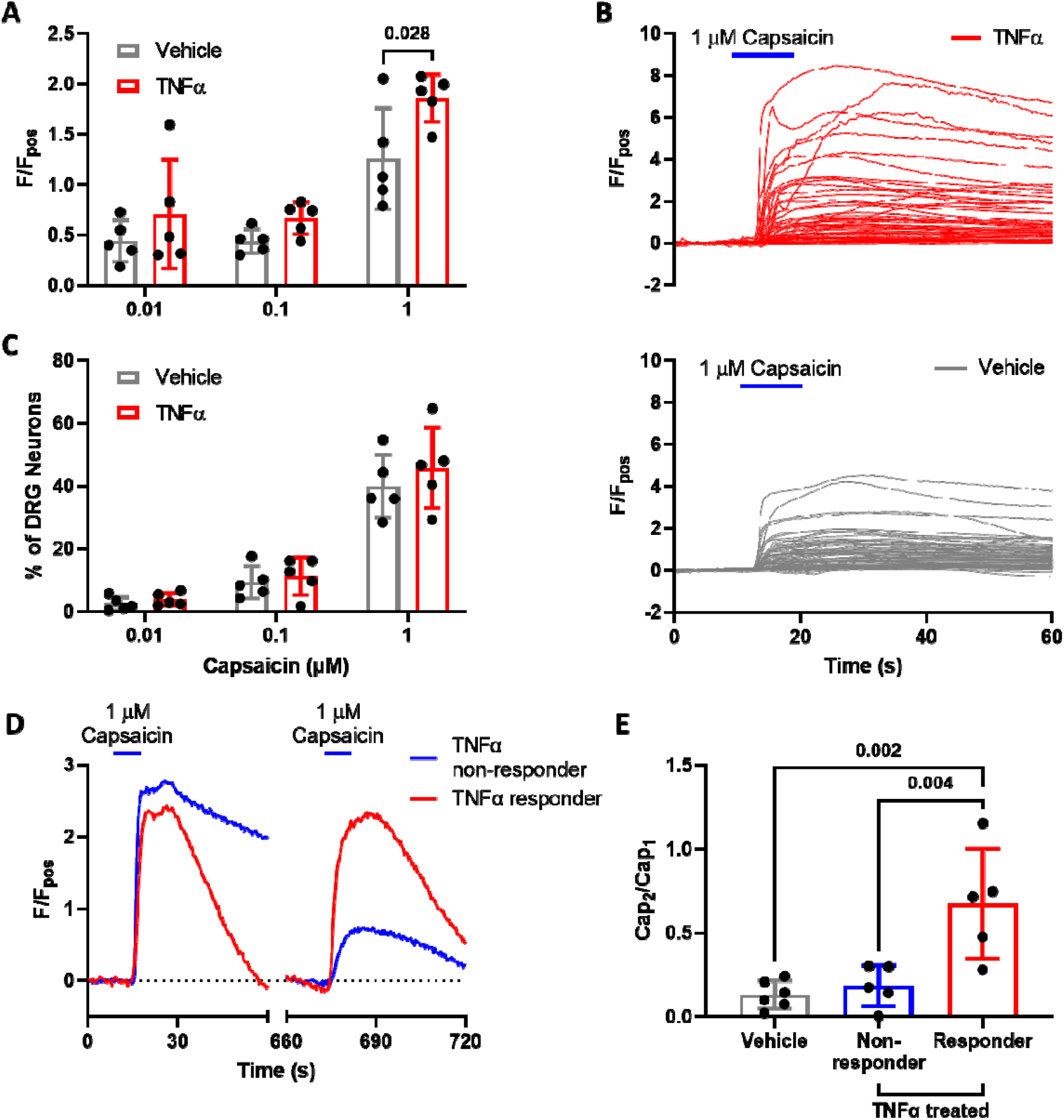
TNFα sensitised capsaicin-evoked [Ca^2+^]_i_ increase in DRG neurons. Capsaicin (1 μM) application resulted in (A) a significantly greater peak response averaged per dish in capsaicin-sensitive DRG neurons pre-incubated with TNFα compared with vehicle (p = 0.028, two-way ANOVA with Holm-Šídák’s multiple comparisons test; n = 5 dishes from N = 5 mice per group). Representative response profiles illustrating (B) the greater effect of 1 μM capsaicin (10 s) on DRG neurons pre-incubated with TNFα (3 nM, top panel) compared with vehicle (PBS, bottom panel) for 24 hours. (C) The overall proportion of capsaicin-sensitive DRG neurons was comparable between culture dishes pre-incubated with vehicle or TNFα respectively at all capsaicin concentrations (p = 0.775, two-way ANOVA with Holm-Šídák’s multiple comparisons test; n = 5, N = 5). In addition, the marked desensitisation of responses to a repeat capsaicin application was greatly attenuated by TNFα, as illustrated (D) by the desensitising response profile to repeat capsaicin application in a TNFα-insensitive (blue) DRG neuron in comparison to the lack of desensitisation to capsaicin in a TNFα-sensitive (red) neuron; and confirmed in (E) by the significantly greater peak response ratios between the second (cap_2_) and first (cap_1_) capsaicin application in DRG neurons co-sensitive to TNFα (3 nM) compared with vehicle (ECS; p = 0.002) or TNFα insensitive neurons (p = 0.004, one-way ANOVA with Holm-Šídák’s multiple comparisons test; n = 5-6 dishes from N = 5-6 independent cultures).

### TNFR1, TRPV1 and p38 MAPK signalling mediates the sensitisation of DRG neurons by TNFα

Having confirmed that TNFα sensitises responses to capsaicin in DRG neurons, we next investigated the rise in [Ca^2+^]_i_ elicited by 1 min TNFα alone. Application of TNFα elicited a concentration-dependent increase in the magnitude of response (p = 0.050, one-way ANOVA with Holm-Šídák’s multiple comparisons test; n = 5, N = 5; Figure 2A & B). Lower concentrations of TNFα did not elicit significantly different response magnitudes (0.03 nM vs 0.1 nM: p = 0.321; 0.1 nM vs 3 nM: p = 0.321, one-way ANOVA with Holm-Šídák’s multiple comparisons test; n = 5, N = 5; Figure 2A & B). TNFα activated a greater proportion of neurons at 3 nM compared to 0.03 nM (p = 0.014, Kruskal-Wallis test with Dunn’s multiple comparisons test; n = 5, N = 5; Figure 2C). No difference in the proportion of TNFα-sensitive neurons was found at other concentrations (0.03 nM vs 0.1 nM: p > 0.999; 0.1 nM vs 3 nM: p = 0.121, Kruskal-Wallis test with Dunn’s multiple comparisons test; n = 5, N = 5; Figure 2C). At the highest capsaicin concentration tested, 45.8 ± 9.37% of DRG neurons were activated by TNFα, of which 79.9 ± 10.7% were also co-sensitive for capsaicin, indicating a preferential activation of nociceptors by TNFα. In DRG neurons isolated from TNFR1^-/-^ mice, the magnitudes of TNFα responses were significantly reduced (p = 0.012, one-way ANOVA with Holm-Šídák’s multiple comparisons test, n = 5-6, N = 5-6; Figure 2D & E) and the change in [Ca^2+^]_i_ in response to TNFα was no different to that produced by ECS (indicated by dashed line in Figure 2F; p = 0.0001, one-way ANOVA with Holm-Šídák’s multiple comparisons test; n = 5-6, N = 5-6). Pre-treatment with R7050, an inhibitor of TNFR1 signalling (Cheng *et al*., 2021), attenuated the TNFα-mediated rise in [Ca^2+^]_i_ in DRG neurons isolated from wild type mice (p = 0.035, one-way ANOVA with Holm-Šídák’s multiple comparisons test; n = 5, N = 5; Figure 2E) and reduced the proportion of TNFα-sensitive neurons (p = 0.004, one-way ANOVA with Holm-Šídák’s multiple comparisons test; n = 5, N = 5; Figure 2F). These observations confirm an essential role for TNFR1 in TNFα-mediated nociceptor stimulation.

**Figure 2.**
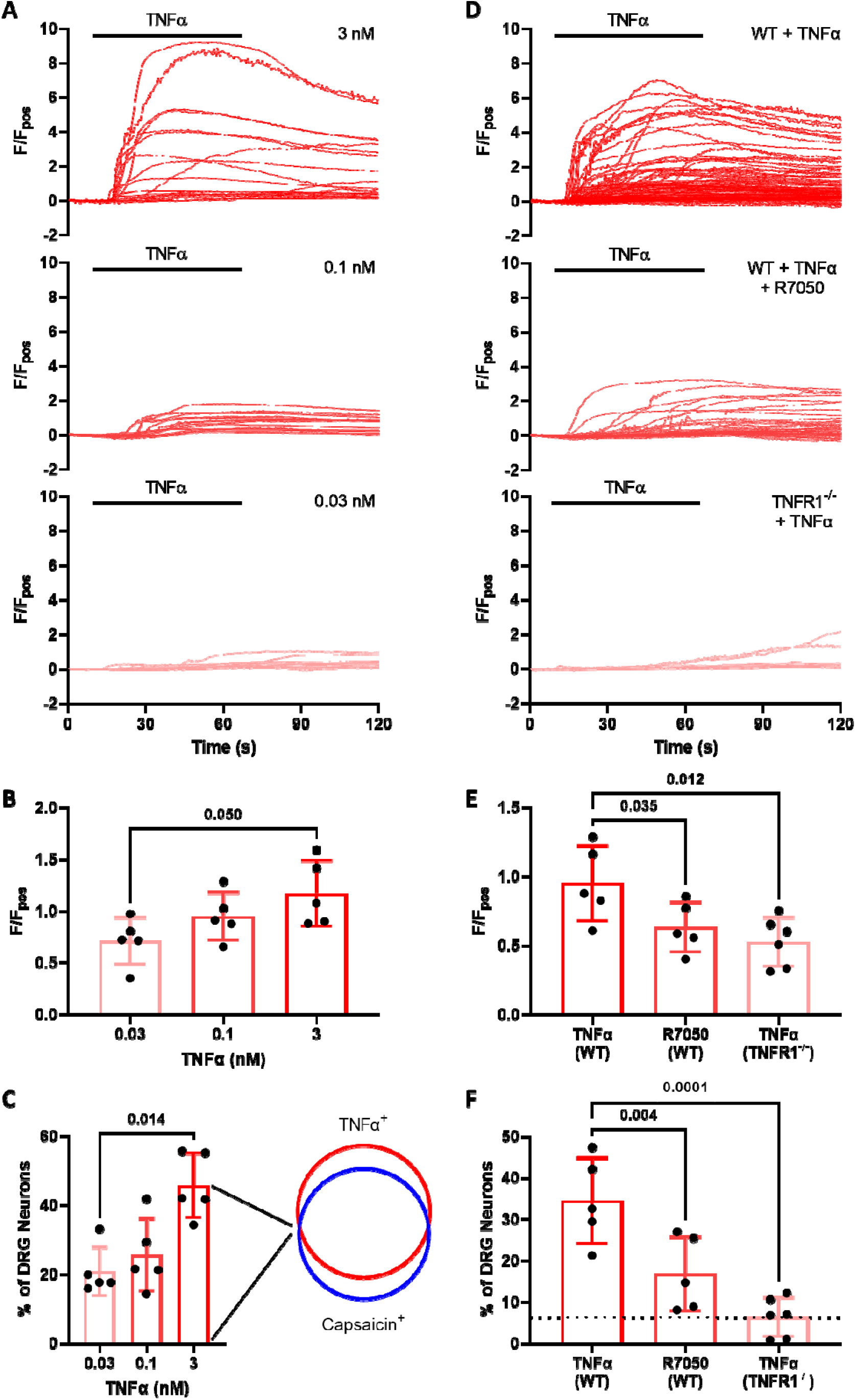
Characterisation of TNFα-evoked [Ca^2+^]_i_ increase in DRG neurons. (A) Representative traces illustrating the concentration-dependent increase in the magnitude of the normalised fluorescent response to TNFα (0.03 - 3.0 nM) within individual neurons from respective culture dishes. (B) The mean magnitude of TNFα responses per dish increased across TNFα concentrations (p = 0.050, one-way ANOVA with Holm-Šídák’s multiple comparisons test; n = 5 dishes from N = 5 independent cultures). (C) The concentration-dependent increase in the proportion (per dish) of TNFα responsive DRG neurons (p = 0.014, Kruskal-Wallis test with Dunn’s multiple comparisons test; n = 5 dishes from N = 5 independent cultures), of which the majority were also co-sensitive to capsaicin. (D) The effects of TNFα (0.1nM) were TNFR1-mediated as illustrated by the significant reduction in (E) response magnitude (p = 0.035; p = 0.012, one-way ANOVA with Holm-Šídák’s multiple comparisons test, n = 5-6 dishes from N = 5-6 independent cultures) and (F) proportion of TNFα-responsive neurons following TNFR1 inhibition with 10 μM R7050 (p = 0.004, one-way ANOVA with Holm-Šídák’s multiple comparisons test; n = 5-6 dishes from N = 5-6 independent cultures) or genetic deletion of TNFR1 in Tnfrsf1a^-/-^ mice in comparison to neurons from wild type (WT) animals (p = 0.0001, one-way ANOVA with Holm-Šídák’s multiple comparisons test; n = 5-6 dishes from N = 5-6 independent cultures). Dotted line represents the proportion of neurons activated in ECS controls (5.54 ± 5.01%, n = 6, N = 6).

Further experiments revealed that the TNFα-mediated increase in [Ca^2+^]_i_ was lost following removal of extracellular Ca^2+^ (p = 0.015, Kruskal-Wallis test with Dunn’s multiple comparisons test; n = 5-8, N = 5; Figure 3A), but unaffected by depletion of intracellular Ca^2+^ stores (p > 0.999, Kruskal-Wallis test with Dunn’s multiple comparisons test; n = 5-8, N = 5), demonstrating that the rise in [Ca^2+^]_i_ was driven by external Ca^2+^ entry. This was further shown by a decrease in TNFα response magnitude following external Ca^2+^ depletion (p = 0.001, Kruskal-Wallis test with Dunn’s multiple comparisons test; n = 5-8, N = 5; Figure 3B), but no change in thapsigargin-treated neurons (p = 0.310, Kruskal-Wallis test with Dunn’s multiple comparisons test; n = 5-8, N = 5). Consistent with this observation, pre-treatment with the TRPV1 antagonist A425619 (1 μM; (Zhang *et al*., 2011)) significantly attenuated the proportion of neurons responding to TNFα (p = 0.037, one-way ANOVA with Holm-Šídák’s multiple comparisons test; n = 5, N = 5; Figure 3C) at a concentration that abolished responses to capsaicin (e.g. 4.39 ± 2.29% in the presence of A425619 vs 44.63 ± 8.19% in the absence of A425619; p < 0.0001, unpaired t-test; n = 4-5, N = 4-5; data not shown). However, A425619 had no effect on the magnitude of responses in the population of neurons still activated by TNFα (p = 0.431, one-way ANOVA with Holm-Šídák’s multiple comparisons test; n = 5, N = 5; Figure 3D). Furthermore, inhibition of TRPA1 with 1 μM AM0902 (Huang *et al*., 2019) also reduced the proportion of TNFα responders (p = 0.001, one-way ANOVA with Holm-Šídák’s multiple comparisons test; n = 5, N = 5; Figure 3C) and the magnitude of TNFα responses compared to controls (p = 0.002, one-way ANOVA with Holm-Šídák’s multiple comparisons test; n = 5, N = 5; Figure 3D, as well as attenuating TNFα Ca^2+^ responses more effectively than A425619 (p = 0.019, one-way ANOVA with Holm-Šídák’s multiple comparisons test; n = 5, N = 5). Co-administration of A425619 and AM0902 to simultaneously inhibit TRPV1 and TRPA1 depleted the number of TNFα-responsive neurons (p < 0.0001, one-way ANOVA with Holm-Šídák’s multiple comparisons test; n = 5, N = 5; Figure 3C) and decreased response magnitude compared to controls (p = 0.003, one-way ANOVA with Holm-Šídák’s multiple comparisons test; n = 5, N = 5; Figure 3D). A combination of A425619 and AM0902 resulted in a smaller proportion of TNFα sensitive neurones compared to use of A425619 (p = 0.027, one-way ANOVA with Holm-Šídák’s multiple comparisons test; n = 5, N = 5) and reduced TNFα responses (p = 0.023, one-way ANOVA with Holm-Šídák’s multiple comparisons test; n = 5, N = 5). These results indicate a TRP-dependent component in TNFα-mediated neuronal activation.

**Figure 3.**
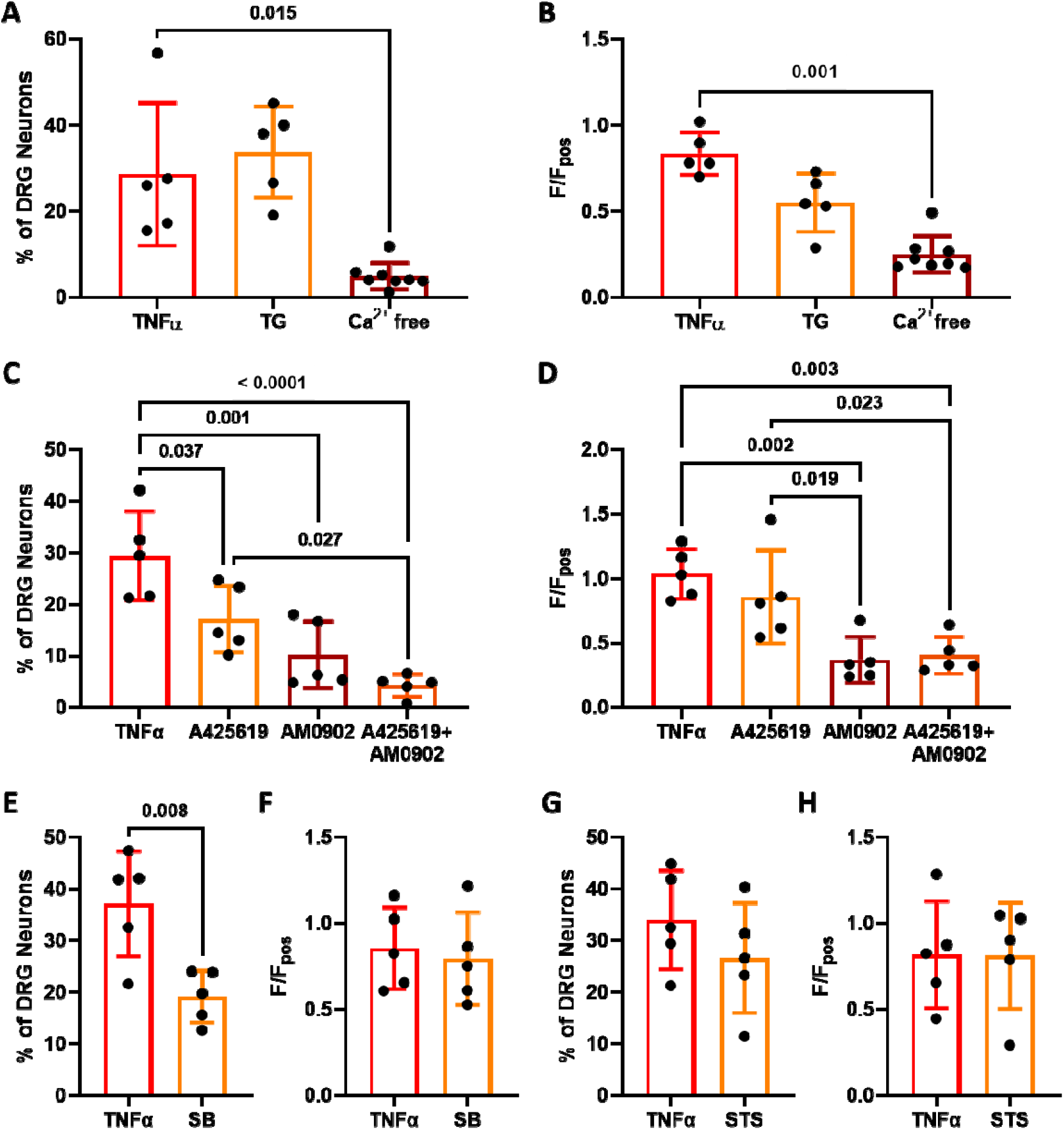
TNFα-evoked [Ca^2+^]_i_ increase is mediated by extracellular Ca^2+^, TRPV1 and p38 MAPK activation. (A) The proportion of DRG neurons responding to 0.1 nM TNFα was unchanged following depletion of internal Ca^2+^ stores with 1 μM thapsigargin (TG; p > 0.999, Kruskal-Wallis test with Dunn’s multiple comparisons test; n = 5-8 dishes from N = 5 independent cultures) and abolished by removal of external Ca^2+^ (p = 0.015, Kruskal-Wallis test with Dunn’s multiple comparisons test; n = 5-8 dishes from N = 5 independent cultures). (B) Depletion of external Ca^2+^ also decreased TNFα response magnitude (p = 0.001, Kruskal-Wallis test with Dunn’s multiple comparisons test; n = 5-8 dishes from N = 5 independent cultures), but was unchanged by TG (p = 0.310, Kruskal-Wallis test with Dunn’s multiple comparisons test; n = 5-8 dishes from N = 5 independent cultures). This effect was partially mediated by TRPV1 and TRPA1 channels, as illustrated (C) by the additive reduction in TNFα-sensitive neurons following pre-incubation with TRPV1 inhibitor 1 μM A425619 and TRPA1 inhibitor 1 μM AM0902 (p < 0.0001, one-way ANOVA with Holm-Šídák’s multiple comparisons test; n = 5 dishes from N = 5 independent cultures), and (D) the reduction in Ca^2+^ responses to TNFα (p 0.003, one-way ANOVA with Holm-Šídák’s multiple comparisons test; n = 5 dishes from N = 5 independent cultures). Furthermore, (E) the proportion of neurons activated by TNFα was significantly attenuated following p38 MAPK inhibition with 10 μM SB203580 (SB; p = 0.008, unpaired t-test; n = 5 dishes from N = 5 independent cultures), but response magnitude was unaffected (p = 0.716, unpaired t-test; n = 5 dishes from N = 5 independent cultures). In contrast, inhibition of protein kinase C (PKC) activity with 10 μM staurosporine (STS) had no effect on (G) the proportion of TNFα responders (p = 0.283, unpaired t-test; n = 5 dishes from N = 5 independent cultures) or (G) the magnitude of TNFα responses (p = 0.977, unpaired t-test; n = 5 dishes from N = 5 independent cultures).

In addition, pre-treatment with the p38 MAPK inhibitor SB203580 (10 μM; (Wu *et al*., 2016)), significantly reduced the proportion of TNFα-sensitive neurons (p = 0.008, unpaired t-test; n = 5, N = 5; Figure 3E) in agreement with previous findings demonstrating a role for p38 MAPK in TNFα-mediated neuronal sensitisation (Gudes *et al*., 2015). SB203580 had no significant effect on TNFα response magnitude (p = 0.716, unpaired t-test; n = 5, N = 5; Figure 3F). In contrast, the protein kinase C (PKC) inhibitor staurosporine (10 μM; (Rusin & Moises, 1998)) failed to reduce the proportion of TNFα-sensitive neurons (p = 0.283, unpaired t-test; n = 5, N = 5; Figure 3G) or response magnitude (p = 0.977, unpaired t-test; n = 5, N = 5; Figure 3H).

Furthermore, in ultrapure DRG neuron cultures, in which non-neuronal cells were removed by MACS (p < 0.0001, unpaired t-test; n = 3, N = 3; Figure 4A & B), the response to TNFα (Figure 4C) was still observed in a comparable proportion of neurons (p = 0.491, unpaired t-test; n = 4-5, N = 4-5; Figure 4D), and the magnitude of TNFα responses was unchanged (p = 0.426, unpaired t-test; n = 4-5, N = 4-5; Figure 4E), thereby confirming that TNFα can directly stimulate sensory neurons, consistent with reported TNFR1 expression in DRG neurons.

**Figure 4.**
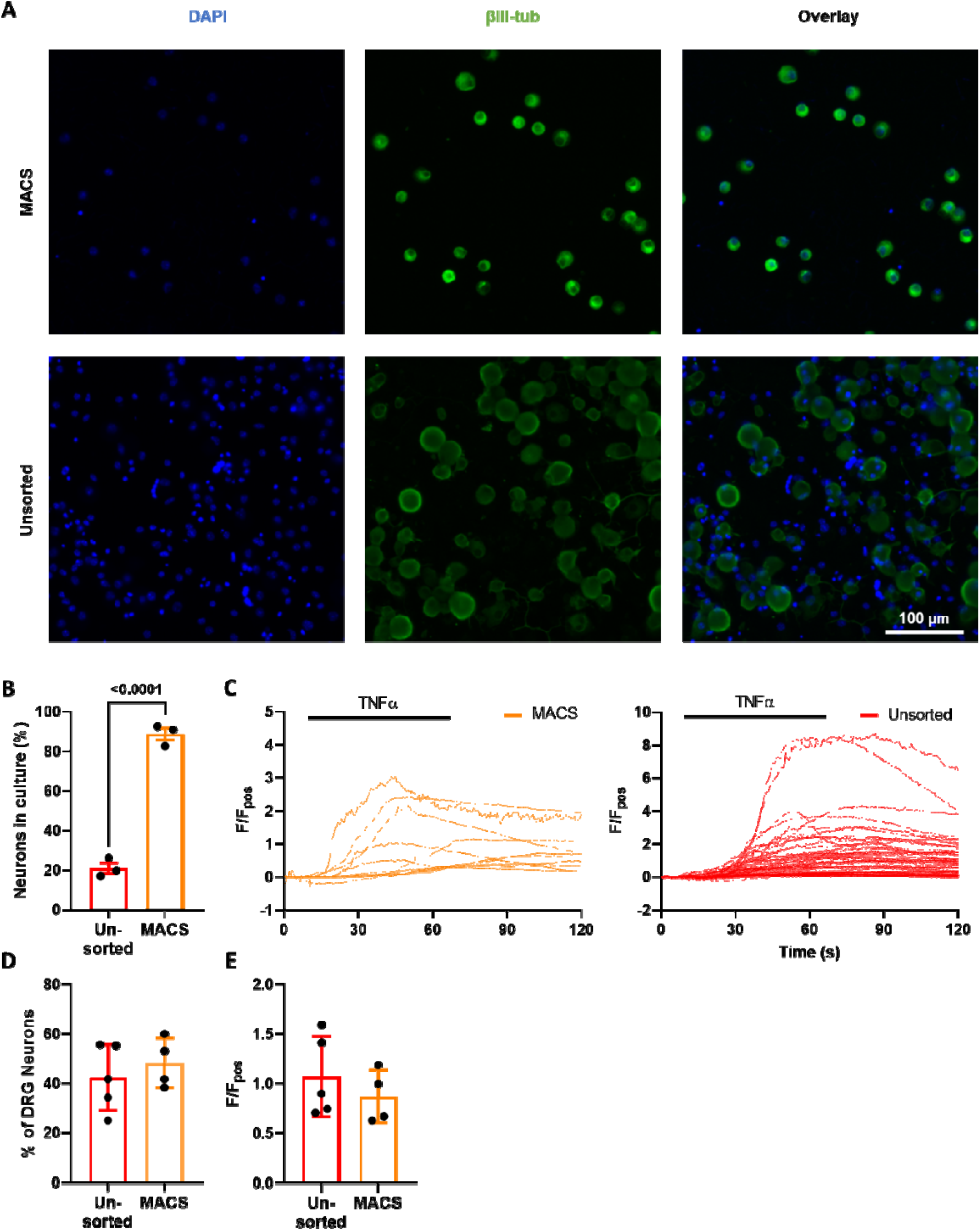
TNFα acts directly on DRG neurons. Magnetic-activated cell sorting (MACS) was used to generate ultra-pure neuronal cultures as illustrated by (A) the immunofluorescent staining of neuronal marker βIII-tubulin (green) and nuclear DAPI stain (blue) in respective cultures and (B) the greatly increased proportion of cells stained with βIII-tubulin following MACS sorting (p < 0.0001, unpaired t-test; n = 3 dishes from N = 3 pooled cultures). TNFα mediated robust increases in [Ca^2+^]_i_ in ultra-pure and unsorted cultures illustrated (C) by individual traces from respective cultures and confirmed by (D) the comparable percentage of TNFα responders in unsorted and MACS sorted DRG cultures (p = 0.491, unpaired t-test; n = 4-5 dishes from N = 4-5 pooled cultures) and (E) equivocal peak responses per dish (p = 0.426, unpaired t-test; n = 4-5 dishes from N = 4-5 pooled cultures).

Having established the important contribution of p38 MAPK signalling and TNFR1 expression to TNFα-mediated Ca^2+^ flux, we next confirmed the involvement of this pathway in the sensitisation of TRPV1 signalling by TNFα. No sensitisation of the magnitude of the [Ca^2+^]_i_ response to capsaicin following 24-hour incubation with TNFα was observed in tissue from TNFR1^-/-^ mice (p = 0.787, unpaired t-test; n = 6, N = 6; Figure 5A & B), and the proportion of capsaicin-sensitive DRG neurons was comparable to controls (p = 0.891, unpaired t-test; n = 6, N = 6; Figure 5C). Following co-incubation of TNFα with SB203580, TNFα-mediated sensitisation of capsaicin responses was attenuated compared to vehicle and SB203580 controls (p = 0.910; p = 0.944, one-way ANOVA with Holm-Šídák’s multiple comparisons test; n = 5, N = 5; Figure 5D & E), and the proportion of responders unchanged (p = 0.380; p = 0.902, one-way ANOVA with Holm-Šídák’s multiple comparisons test; n = 5, N = 5; Figure 5F).

**Figure 5.**
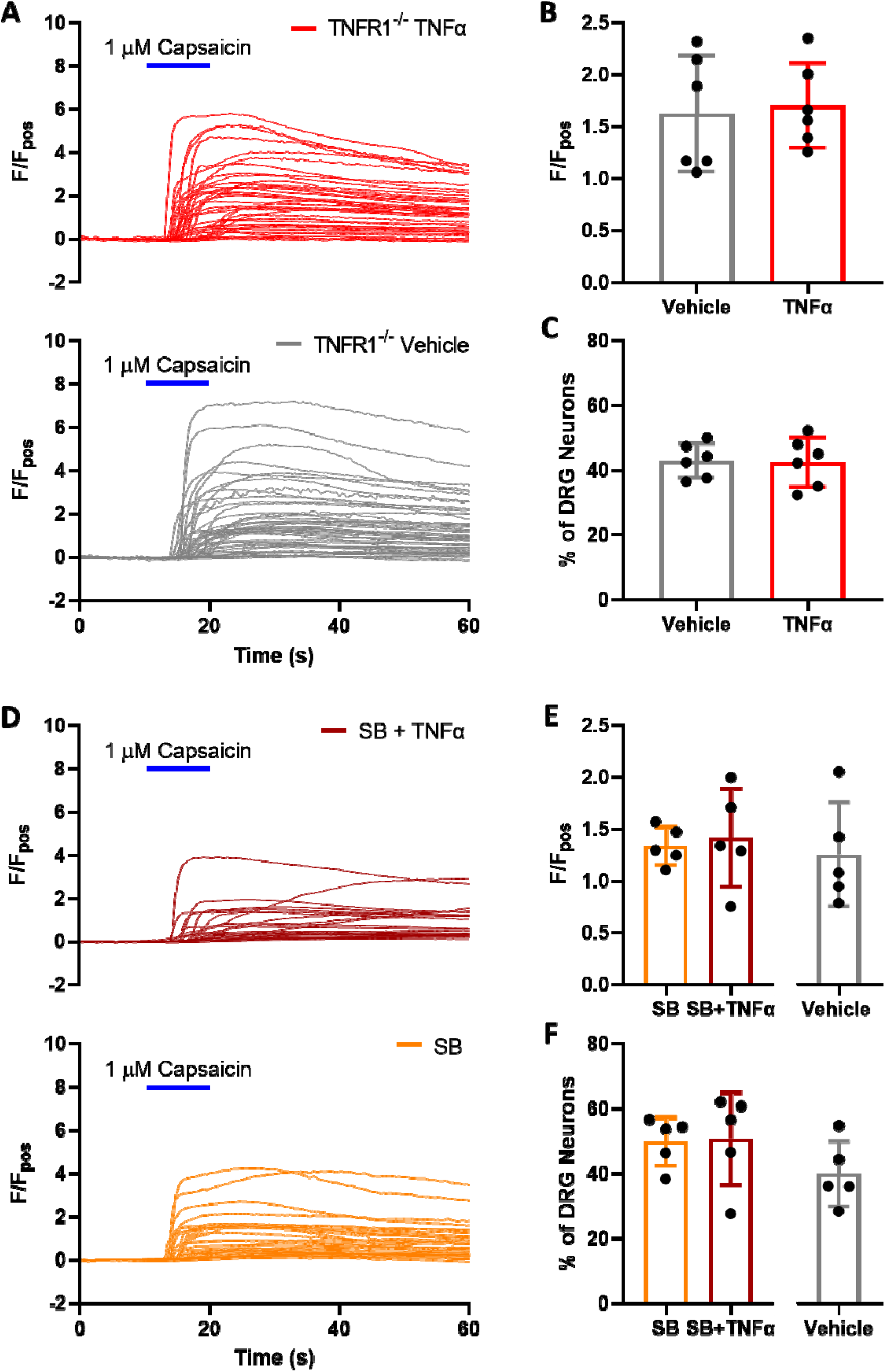
TNFα sensitisation of capsaicin-evoked [Ca^2+^]_i_ increase in DRG neurons is p38 MAPK-and TNFR1-mediated. (A) Example traces of individual response profiles to 1 μM capsaicin (10 s) in DRG neurons cultured from TNFR1^-/-^ mice following 24-hour treatment of respective culture dishes with vehicle (PBS) or TNFα (3 nM). (B) Incubation with TNFα no longer increased peak averaged per dish Ca^2+^ responses to capsaicin in DRG neurons from TNFR1^-/-^ mice (p = 0.787, unpaired t-test; n = 6 dishes from N = 6 independent cultures). (C) The proportion of capsaicin-sensitive neurons remained comparable between culture dishes pre-incubated with vehicle or TNFα in DRG neurons from TNFR1^-/-^ mice (p = 0.891, unpaired t-test; n = 6 dishes from N = 6 independent cultures). Similarly, overnight incubation with TNFα no longer increased the magnitude of capsaicin responses in DRG neurons co-incubated with the p38 MAPK inhibitor SB203580 as illustrated by: (D) individual neuronal responses to 1 μM capsaicin (10 s) in SB203580-treated DRG neurons following 24h incubation with vehicle or TNFα (3 nM); (E) the comparable magnitude of the peak averaged per dish response to capsaicin between respective treatments (p = 0.944, one-way ANOVA with Holm-Šídák’s multiple comparisons test; n = 5 dishes from N = 5 independent cultures). (F) The proportion of capsaicin-sensitive DRG neurons was also comparable between treatment groups following p38 MAPK inhibition (p = 0.380, one-way ANOVA with Holm-Šídák’s multiple comparisons test; n = 5 dishes from N = 5 independent cultures).

### TNFα sensitises colonic afferent responses to noxious ramp distension and capsaicin via TNFR1 and p38 MAPK

Finally, to confirm the translation of our findings from DRG neurons to the activation of colonic afferents, we studied the contribution of TNFR1 and p38 MAPK signalling to TNFα-mediated sensitisation of colonic afferent responses to noxious ramp distension and capsaicin (Figure 6A). Consistent with the sensitising effects of TNFα observed previously in DRG neurons, treatment with TNFα prevented the desensitisation of colonic afferent responses to repeated ramp distensions compared to vehicle (p = 0.0004, two-way ANOVA with Holm-Šídák’s multiple comparisons test; N = 8; Figure 6B). Comparisons within treatment groups demonstrated a significant increase in afferent response between ramp distensions 3 and 5 in TNFα-treated tissues (p = 0.011, two-way ANOVA with Holm-Šídák’s multiple comparisons test; N = 8), whereas no significant change occurred between responses to ramp distensions 3 and 5 in vehicle-treated tissues (p = 0.425, two-way ANOVA with Holm-Šídák’s multiple comparisons test; N = 8). Significantly greater afferent responses to ramp distension were observed across noxious distending pressures (p = 0.0002, multiple unpaired t-tests, N = 8; Figure 6C) and peak afferent firing was significantly increased (p = 0.0004, unpaired t-test, N = 8; Figure 6D). An enhanced afferent response to capsaicin was also observed following application of TNFα compared with vehicle (p = 0.003, Mann-Whitney test; N = 8; Figure 7A & B). TNFα significantly increased nerve firing in the 10 minutes post capsaicin application (p = 0.005, two-way ANOVA with Holm-Šídák’s multiple comparisons test; N = 8; Figure 7C) and peak activity was elevated 75% compared to vehicle (p = 0.007, unpaired t-test, N = 8; Figure 7D). Compliance of distensions was comparable between treatment groups (e.g., 0.741 ± 0.137 ml vs 0.731 ± 0.074 ml for the 5^th^ ramp distension in TNFα vs vehicle treated tissues, p = 0.842, unpaired t-test; N = 8; data not shown).

**Figure 6.**
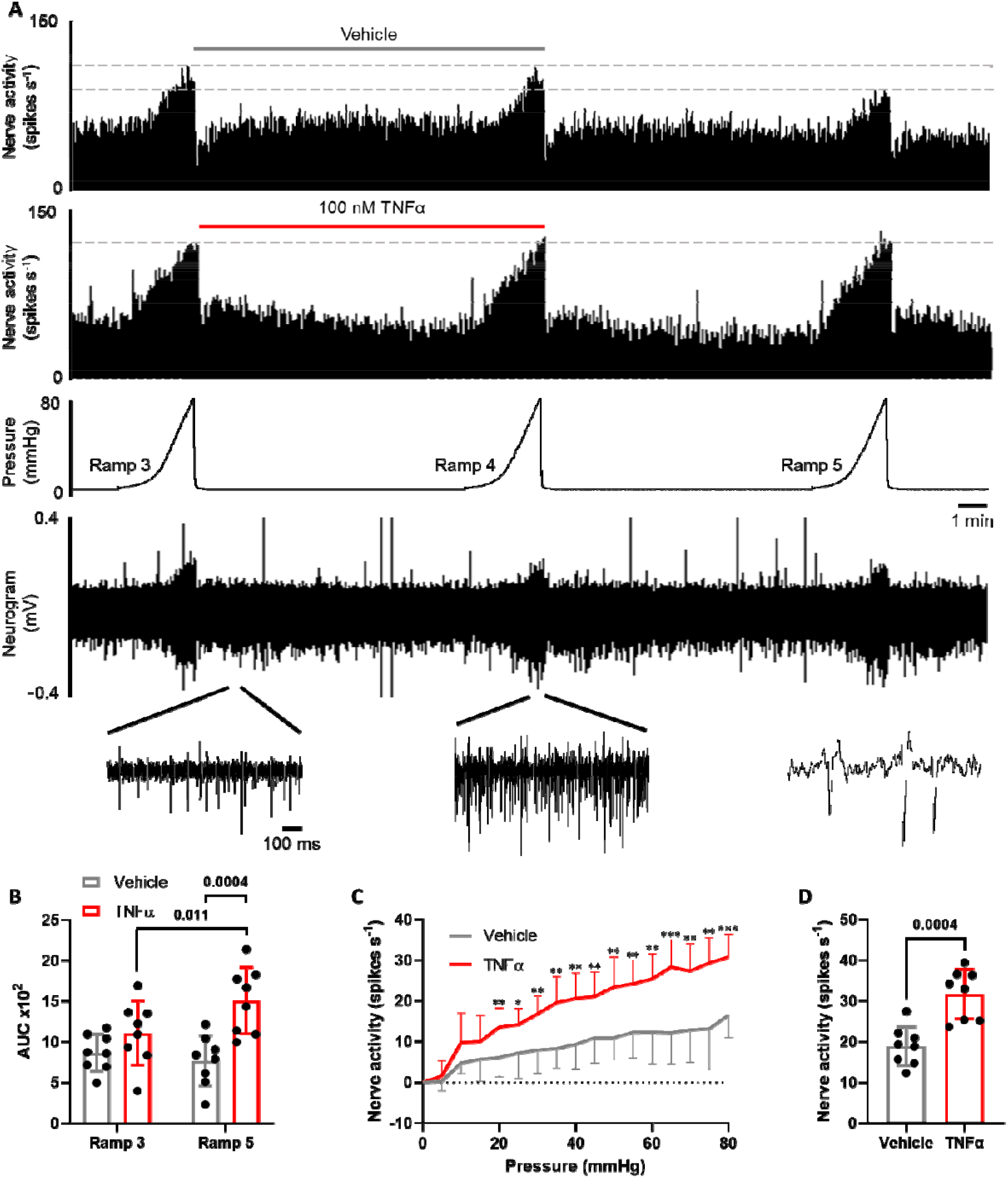
TNFα evokes colonic afferent mechanical hypersensitivity. (A) Example rate histograms and neurogram of lumbar splanchnic nerve (LSN) activity with accompanying pressure trace showing sequential (x3) ramp distensions (0-80 mmHg) from vehicle- and TNFα (100 nM)-treated preparations, highlighting the desensitisation of afferent responses to distension no longer occurred following TNFα treatment. This effect was confirmed by: (B) the significantly greater afferent response to ramp distension (measured by area under the curve, AUC, during ramp distension) following TNFα pre-treatment compared with vehicle (ramp 5) (p = 0.0004, two-way ANOVA with Holm-Šídák’s multiple comparisons test; N = 8) and within treatment groups (p = 0.011; two-way ANOVA with Holm-Šídák’s multiple comparisons test; N = 8); (C) the significantly greater increase in afferent response throughout the noxious distending pressure range (< 20mmHg, ramp 5) following TNFα treatment compared with vehicle (* p < 0.05, ** p < 0.01, *** p < 0.001, multiple t-tests; N = 8 animals); (D) the significantly greater peak increase in afferent discharge to ramp distension following application of TNFα compared with vehicle (p = 0.0004, unpaired t-test; N = 8).

**Figure 7.**
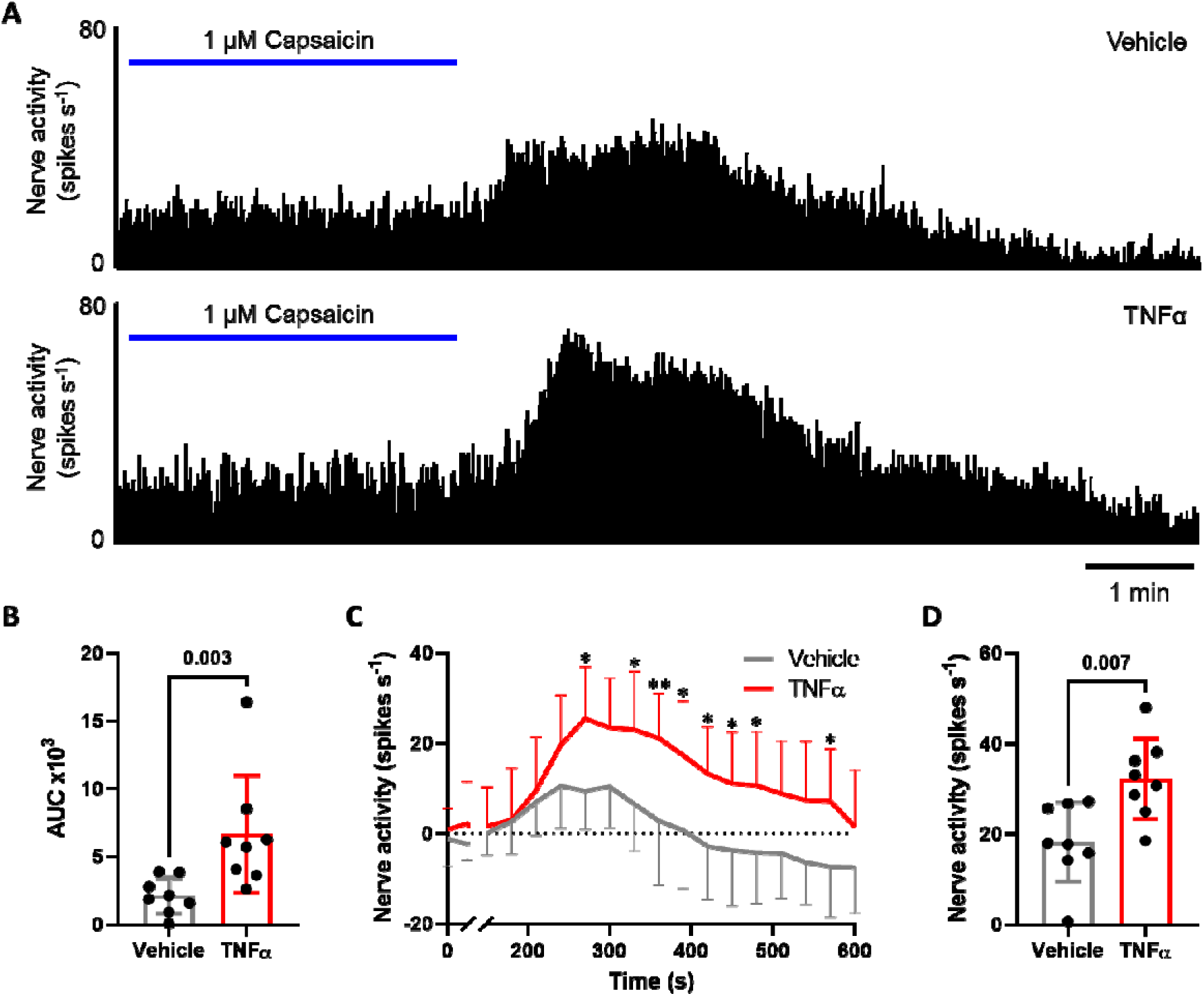
TNFα sensitises colonic afferent responses to capsaicin. (A) Example rate histograms of LSN recordings illustrating the greater afferent response to 1 μM capsaicin following TNFα (100 nM) compared with vehicle pre-treatment. This was confirmed by the significantly greater afferent discharge following TNFα compared with vehicle pre-treatment measured by: (B) AUC of afferent firing 0-10 min post capsaicin application (p = 0.003, Mann-Whitney test; N = 8 animals); (C) prolonged increase in afferent discharge to capsaicin (* p < 0.05, ** p < 0.01, two-way ANOVA with Holm-Šídák’s multiple comparisons test; N = 8 animals); (D) greater peak change in afferent firing 0-10 min after application of capsaicin (p = 0.007, unpaired t-test; N = 8 animals).

In keeping with data from DRG neurons, where we observed a reliance on TNFR1 for the sensitising effects of TNFα, no sensitisation of responses to noxious ramp distension was observed following the application of TNFα compared with vehicle in tissue from TNFR1^-/-^ mice (p = 0.521, unpaired t-test; N = 7-8; Figure 8A). In addition, no difference in afferent activity was seen across noxious distending pressures (p = 0.280, multiple unpaired t-tests; N = 7-8; Figure 8B) and peak afferent firing was unchanged between vehicle- and TNFα-treated TNFR1^-/-^ tissue (p = 0.666, unpaired t-test; N = 7-8; Figure 8C). No TNFα-mediated sensitisation was observed in response to capsaicin (p = 0.767, unpaired t-test; N = 7-8; Figure 8D), and afferent firing was comparable 10 minutes after capsaicin application (p = 0.961, two-way ANOVA with Holm-Šídák’s multiple comparisons test; N = 7-8; Figure 8E). Furthermore, TNFα had no effect on peak afferent activity in the absence of TNFR1 (p = 0.967, unpaired t-test; N = 7-8; Figure 8F). Interestingly, responses to ramp distension (AUC: WT 769 ± 312 vs TNFR1^-/-^ 1665 ± 412; p = 0.0004, unpaired t-test; N = 7-8) and capsaicin (AUC: WT 2114 ± 1335 vs TNFR1^-/-^ 5615 ± 2535; p = 0.005, unpaired t-test; N = 7-8) were significantly higher in vehicle treated tissues from TNFR1^-/-^ mice compared to wild type (WT) mice.

**Figure 8.**
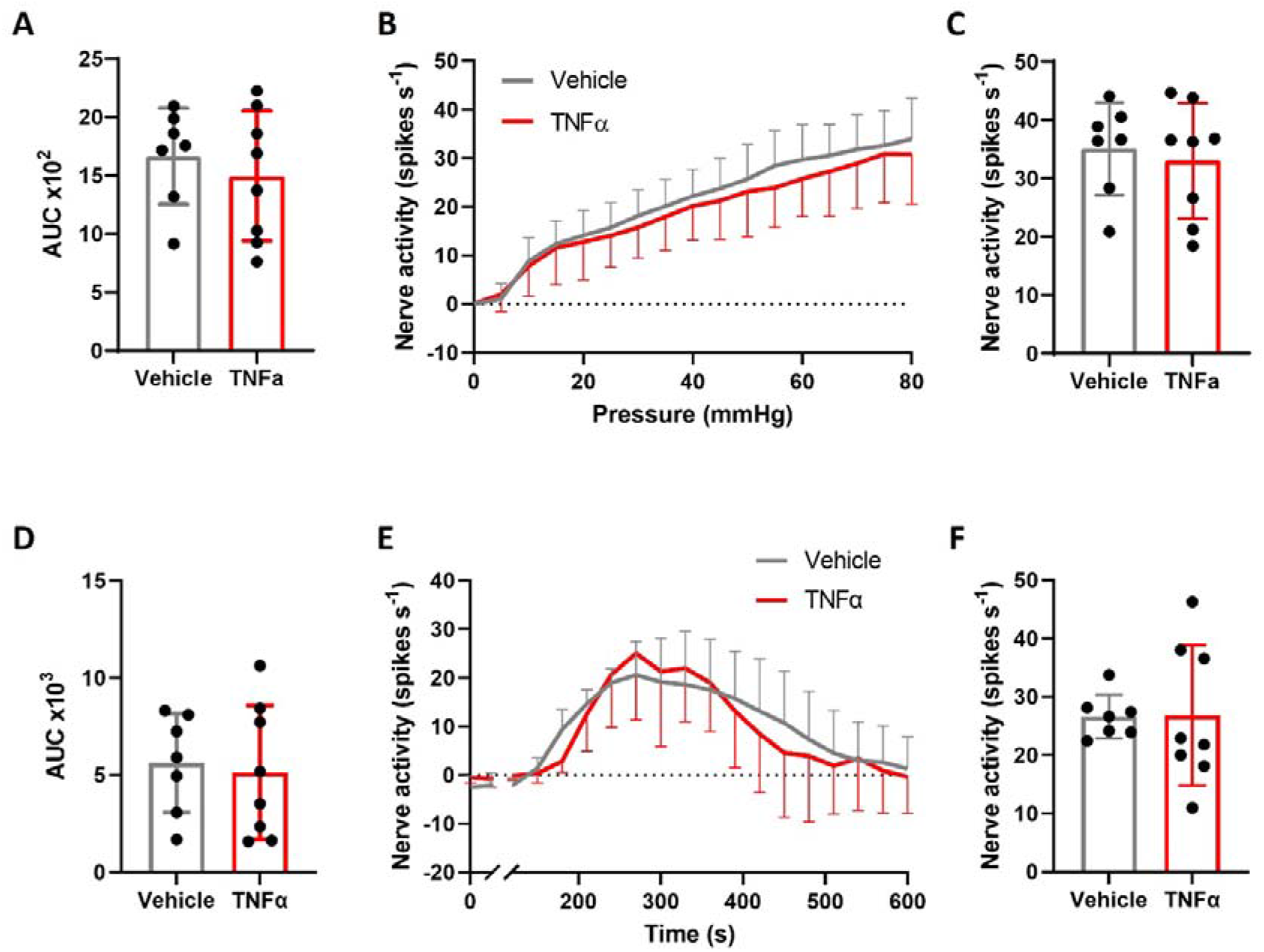
TNFR1 mediates TNFα-induced colonic afferent sensitisation. The sensitisation of colonic afferents by TNFα pre-treatment was dependent on TNFR1 expression as confirmed by the presence of comparable colonic afferent responses to ramp distension and capsaicin following vehicle or TNFα pre-treatment in tissue from TNFR1^-/-^ mice shown by: (A) comparing the AUC of nerve activity following ramp distension (p = 0.521, unpaired t-test; N = 7-8); (B) afferent response profiles to ramp distension (p = 0.280, multiple t-tests; N = 7-8 animals); (C) peak firing frequency to ramp distension (p = 0.666, unpaired t-test; N = 7-8); (D) AUC of firing frequency following capsaicin application (p = 0.767, unpaired t-test; N = 7-8); (E) afferent response profiles to capsaicin (p = 0.961, two-way ANOVA with Holm-Šídák’s multiple comparisons test; N = 7-8 animals); (F) the peak increase in afferent discharge to capsaicin (p = 0.967, unpaired t-tests; N = 7-8).

In agreement with our identification of a role for p38 MAPK signalling in the sensitising effects of TNFα in DRG neurons, we observed no sensitisation of colonic afferents to noxious ramp distension following the application of TNFα in the presence of SB203580 to inhibit p38 MAPK (p = 0.042, Kruskal-Wallis test with Dunn’s multiple comparisons test; N = 6-8; Figure 9A) or in SB203580 controls (p = 0.011, Kruskal-Wallis test with Dunn’s multiple comparisons test; N = 6-8). SB203580 attenuated TNFα sensitisation across all noxious distending pressures (p = 0.004; multiple unpaired t-tests; N = 7-8; Figure 9B) and p38 MAPK inhibition significantly decreased peak afferent firing following TNFα incubation (p = 0.026, Kruskal-Wallis test with Dunn’s multiple comparisons test; N = 6-8; Figure 9C). As expected, no sensitisation was observed in SB203580 controls (p = 0.020, Kruskal-Wallis test with Dunn’s multiple comparisons test; N = 6-8; Figure 9C). Furthermore, SB203580 significantly attenuated afferent responses to capsaicin in TNFα-treated tissues (p = 0.034, Kruskal-Wallis test with Dunn’s multiple comparisons test; N = 7-8; Figure 9D), with significantly reduced nerve activity profiles (p = 0.0002, two-way ANOVA with Holm-Šídák’s multiple comparisons test; N = 7-8; Figure 9E). Responses to capsaicin in SB203580 controls were significantly lower than TNFα-treated tissues (p = 0.017, Kruskal-Wallis test with Dunn’s multiple comparisons test; N = 7-8; Figure 9D). Additionally, SB203580 reversed the TNFα-mediated increase in peak afferent firing (p = 0.022, one-way ANOVA with Holm-Šídák’s multiple comparisons test; N = 7-8; Figure 9F) and as expected peak firing was unchanged in SB203580 controls (p = 0.022, one-way ANOVA with Holm-Šídák’s multiple comparisons test; N = 7-8; Figure 9F).

**Figure 9.**
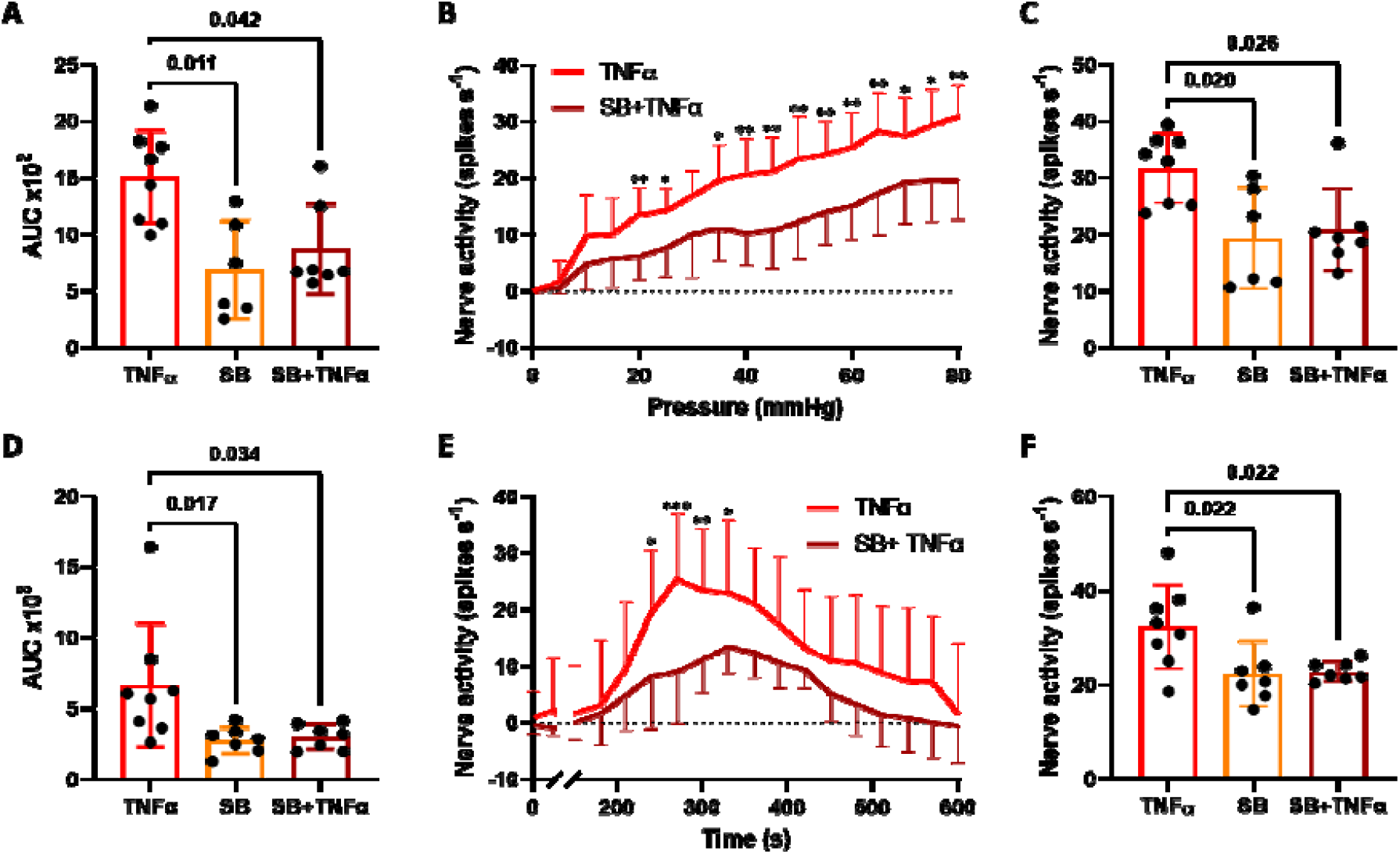
TNFα-induced colonic afferent sensitisation is mediated by p38 MAPK. The ability of TNFα to sensitise colonic afferent responses to ramp distension and capsaicin was abolished by co-administration with the p38 MAPK inhibitor SB203589 as demonstrated by: (A) the significant reduction in AUC of colonic afferent responses to ramp distensions following pre-treatment with TNFα in the presence of SB203589 (p = 0.042, Kruskal-Wallis test with Dunn’s multiple comparisons test; N = 6-8 animals); (B) the significant reduction in afferent responses to ramp distension across noxious distending pressures (< 20mmHg) following pre-treatment with TNFα in the presence of SB203580 (* p < 0.05, ** p < 0.01, multiple t-tests; N = 6-8 animals); (C) the significant reduction in the peak afferent response to ramp distension following application of TNFα in the presence of SB203580 (p = 0.026, Kruskal-Wallis test with Dunn’s multiple comparisons test; N = 6-8 animals); (D) significantly reduced afferent responses to capsaicin measured by AUC following pre-treatment with TNFα in the presence of SB203580 (p = 0.034, Kruskal-Wallis test with Dunn’s multiple comparisons test; N = 7-8 animals); (E) the significant reduction in afferent responses to capsaicin at multiple time points following pre-treatment with TNFα in the presence of SB203580 (* p < 0.05, ** p < 0.01, *** p < 0.001, two-way ANOVA with Holm-Šídák’s multiple comparisons test; N = 7-8 animals); (F) the significant reduction in peak nerve discharge to capsaicin following pre-treatment with TNFα in the presence of SB203580 (p = 0.022, one-way ANOVA Holm-Šídák’s multiple comparisons test; N = 7-8 animals).

## Discussion

TNFα has been linked to the production of abdominal pain in gastrointestinal disease due to its enhanced expression in disease states such as IBS (Hughes *et al*., 2013) and IBD (Kamada *et al*., 2008), combined with data showing TNFα-mediated visceral nociceptor sensitisation and TNFR1 expression in colonic nociceptors (Hockley *et al*., 2019). The signalling pathways mediating these effects have not been fully explored, although data from studies of somatic hypersensitivity point to a role for p38 MAPK signalling downstream of TNFR1 receptor activation (Jin & Gereau IV, 2006) and a sensitising effect on TRPV1, a detector of noxious stimuli (Khan *et al*., 2008). As such, the goal of this study was to investigate the contribution of TNFR1 and p38 MAPK signalling to the pro-nociceptive effects of TNFα including TRPV1 receptor activation in sensory neurons and colonic afferents.

Data from our studies confirmed that TNFα sensitises TRPV1 receptor signalling, by demonstrating: i) an enhanced capsaicin-evoked increase in [Ca^2+^]_i_ within sensory DRG neurons following overnight incubation with TNFα; ii) a TRPV1 receptor-mediated increase in [Ca^2+^]_i_ to acute application of TNFα in sensory DRG neurons that subsequently showed significantly less desensitisation to repeated application of capsaicin; and iii) a marked increase in the colonic afferent response to capsaicin following TNFα pre-treatment. In keeping with our hypothesis, these effects were no longer observed in tissue from TNFR1^-/-^ mice and were greatly attenuated by pre-treatment with the p38 MAPK inhibitor SB203580; conditions in which TNFα also no longer prevented the desensitisation of colonic afferent responses to repeated noxious distension of the bowel. Collectively these findings provide further functional evidence of a contribution by TNFα to the production of visceral pain in gastrointestinal disease and highlight the role of TNFR1-mediated p38 MAPK signalling to the pro-nociceptive activity of TNFα.

The validity of our data is supported by comparable observations showing TNFα enhances TRPV1 receptor signalling in nodose (Hsu *et al*., 2017; Lin *et al*., 2017), trigeminal (Khan *et al*., 2008) and DRG neurons (Hensellek *et al*., 2007). These effects have been attributed to TNFR1 activation and p38 MAPK signalling, consistent with our data demonstrating the essential contribution of TNFR1 and p38 MAPK to TNFα sensitisation of capsaicin responses in colonic afferents for the first time, and DRG neurons for the first time in the same study. The mechanism by which p38 MAPK modulates TRPV1 receptor activity was not a focus of our experiments, however we observed changes following brief application of TNFα indicative of an acute increase in channel activity that would be expected following phosphorylation of TRPV1 by p38 MAPK. This observation was in keeping with previous studies showing enhanced TRPV1-mediated inward currents shortly after application of

TNFα; effects reported to be mediated by p38 MAPK and protein kinase C (PKC) (Constantin *et al*., 2008). However in contrast to p38 MAPK inhibition, we found no change in the TRPV1-mediated increase in [Ca^2+^]_i_ to TNFα following PKC inhibition with staurosporine. These findings suggest that phosphorylation sites on intracellular domains of TRPV1, which are not specific to PKC, may be targeted and phosphorylated by p38 MAPK. In addition, p38 MAPK signalling has also been shown to increase TRPV1 channel expression following incubation with TNFα for periods of 1 hour or greater (Constantin *et al*., 2008), and this process may also have contributed to the increased capsaicin response we observed following overnight incubation with TNFα.

Consistent with the reported co-expression of TNFR1 with TRPV1 in sensory DRG neurons (Zeisel *et al*., 2018), we found marked co-sensitivity of TNFα-stimulated DRG neurons to capsaicin. This response was dependent on TNFR1 expression and was preserved in ultra-pure, neuron only DRG cultures demonstrating that TNFα can directly modulate sensory neuron activity and TRPV1 signalling via TNFR1 in the same neuron. Furthermore, these data indicate that TNFα-responsive neurons are predominantly nociceptors based on the ability of capsaicin to evoke significant pain in humans, including visceral pain following colorectal administration (van Wanrooij *et al*., 2014).

This observation was corroborated by our finding that TNFα sensitises colonic afferents at noxious distending pressures. These experiments provide evidence that TNFα causes abdominal pain through TNFR1-mediated colonic nociceptor sensitisation due to p38 MAPK-enhanced TRPV1 signalling.

However, this is not the only mechanism by which TNFα stimulates sensory neurons. For example, the increase in [Ca^2+^]_i_ to TNFα, which is solely dependent on extracellular Ca^2+^ entry, was only partially blocked by the TRPV1 antagonist (A425619) at concentrations that completely abolished the Ca^2+^ response to capsaicin. This indicates that other Ca^2+^ permeable ion channels are stimulated by TNFα and the additive inhibitory effect of the TRPA1 antagonist (AM0902) on TNFα-mediated neuronal activation is consistent with existing data implicating TRPA1 in TNFα-mediated afferent sensitisation (Hughes *et al*., 2013). In addition, we also demonstrated that TNFα-mediated colonic afferent mechanosensitisation was dependent on TNFR1 expression and p38 MAPK activity in keeping with seminal data showing this signalling pathway mediates somatic mechanical hypersensitivity to TNFα (Jin & Gereau IV, 2006). These effects have been attributed to enhanced TTX-resistant Na_V_1.8 currents, an effect also seen in response to TNFR1 dependent TNFα signalling in colon projecting DRG neurons. With the inclusion of our study, current data indicates that TNFα has the capacity to sensitise colonic nociceptors through p38 MAPK-enhanced TRPV1, TRPA1 and Na_V_1.8 channel activity downstream of TNFR1.

These findings highlight the utility of TNFα and its downstream signalling pathways as important drug targets for pain relief in gastrointestinal diseases, such as IBS and IBD, in which enhanced TNFα expression has been reported.

## Additional Information

### Data availability

All data supporting the results presented in the manuscript are included in the manuscript figures and raw data sets are available at DOI: 10.17863/CAM.81112.

### Author Contributions

K.H.B. designed the research studies, conducted the experiments, acquired and analysed the data and wrote the manuscript. J.P.H. acquired and analysed the data and wrote the manuscript. T.S.T. acquired the data. L.A.P. analysed the data. I.P.C. designed the research studies. DCB, E.St.J.S. and FW designed the research studies and wrote the manuscript. All authors approved the final version of the manuscript submitted for publication and agree to be accountable for all aspects of the work in ensuring that questions related to the accuracy or integrity of any part of the work are appropriately investigated and resolved. All persons designated as authors qualify for authorship, and all those who qualify for authorship are listed.

